# Reliable and accurate gene expression quantification with subpopulation structure-aware constraints for single-cell RNA sequencing

**DOI:** 10.1101/2022.11.08.515740

**Authors:** Ching-Chih Tu, Jui-Hung Hung

## Abstract

**Background:** Single-cell RNA sequencing (scRNA-seq) analysis analyzes the type and state of individual cells by estimating the gene expression of each cell and enables researchers to study the biological phenomena that cannot be observed in bulk RNA sequencing.

**Motivation:** However, the current scRNA-seq quantification tools estimate the gene expression profile of each cell independently, ignoring the fact that there are multiple cell types in the scRNA-seq data and the expression level should be highly correlated with the cell type. Since scRNA-seq suffers from a low sequencing depth, the conventional strategy leads to a high proportion of missing values in the gene expression profile, obscuring the biological characteristics of cell subpopulations and further impacting the correctness of the subsequent downstream analysis.

**Results:** In this study, we proposed Quasic, a novel scRNA-seq quantification pipeline which examines the potential cell subpopulation information during quantification, and uses the information to calculate the gene expression level. Using the human peripheral blood mononuclear cells and the simulated doublet dataset, we verified that Quasic not only correctly reinforced the cell signatures, but also identified the corresponding cell subpopulations and biological pathways more accurately. In addition, we also applied Quasic to the breast cancer cell line dataset (MCF-7), and successfully identified more potentially therapeutic resistant cells of which characteristics are consistent with that from previous studies.

**Conclusions:** The proposed pipeline can let the gene expression profile of each cell be more consistent with the corresponding subpopulation, making the biological features unique to the subpopulation more apparent and convenient for analysis. By using Quasic, researchers can effectively extract the desired cell subpopulation information from their sampled cells, enable them to perform cell subpopulation-related studies more accurately.

## Introduction

Single-cell RNA sequencing (scRNA-seq) is a high-throughput technology that can measure the expression of the entire transcriptome of each cell. By analyzing scRNA-seq data, researchers can detect novel biological discoveries omitted by traditional bulk RNA sequencing methods, such as revealing rare and crucial cell populations among cells, displaying regulatory relationships between cell types and genes, or even tracking the trajectories of cell lineages in development.

In recent years, multiple tools have been developed to perform quantification of scRNA-seq, which represents exanimating the gene expression level from the raw reads generated by the sequencer, such as Cell Ranger^1^, Alevin^2^, Kallisto^3^ and STARsolo^4^. These quantification tools use different strategies to handle specific demands. For example, Cell Ranger and STARsolo use a general aligner called STAR^5^ to perform the alignment, while Alevin and Kallisto choose to use pseudo alignment to reduce the time-consuming problems of STAR. Moreover, these four quantification tools also perform different cell barcode correction strategies and use different methods to quantify the unique molecular identifiers (UMI), leading to the estimation of gene expression profiles as well as the downstream analysis (such as finding the differential expression genes) perform diversely to some extent^6^.

Although each of these quantification tools has been wildly used by the researchers of scRNA-seq to date, there are some crucial problems in current quantification tools. First of all, since the information on cell subpopulation within the scRNA-seq data often plays an important role in scRNA-seq downstream analysis, the subpopulation structure in the sampled cells is often ambiguous. More specifically, due to the low sampling rate of scRNA-seq, there may be a high proportion of missing values or biased estimation in scRNA-seq data, which causes the difference of biological features between clusters to be usually inconspicuous. Cells may express dissimilarly with other cells even if they have shared the same cell type label (see **Fig. S1**). Moreover, the scarcity of gene expression in sequenced cells or the whole subpopulation also makes researchers hard to extract the characteristics from them, and the examination of cell subpopulation analysis, which is often the main goal of scRNA-seq analysis, would be hard to perform accurately. Generally, researchers can deal with the above problems using imputation tools to infer the missing values in gene expression profiles based on various statistical models. However, currently developed imputation tools often induce false signals which may mislead the researcher^7^.

To improve the quantification of single-cell RNA sequencing data, in this study, we proposed a subpopulation-aware scRNA-seq quantification pipeline called Quasic (**qua**ntification of **si**ngle cell with subpopulation **s**tructure-aware **c**onstraints), which considers the subpopulation information during the optimization for quantification. Quasic aims to (1) facilitate cell type identification, especially for the cells with low gene expression, (2) strengthen the features of cell subpopulations and make the characteristics captured by downstream analyses more robust and precise, and (3) enhance the difference between cell subpopulations, making the results of cell type comparison more significant. In the following paragraph, we would demonstrate the advantages of subpopulation-aware quantification, and how it reinforces the scRNA-seq downstream analysis for the researchers.

## Results

### Framework overview

To begin with, we first assume that cells in the same biological subpopulation should collectively share the same genetic characteristics and that the characteristics of subpopulations are distinct from each other. To model this during the gene expression estimation, we designed a quantification method called subpopulation-aware quantification, which constrains the gene profile by weighing the gene expression levels calculated by original quantification and the corresponding cluster signature for the cell. Moreover, the weighing would be determined by the cell-specific term that represents the confidence level of the cluster results for a particular cell (see **Methods**).

In addition to the subpopulation-aware quantification, the entire framework could be divided into three stages (see **Fig. 1**), including 1) initial quantification, 2) clustering, and 3) subpopulation-aware quantification. In the initial quantification stage, our framework executes traditional quantification implemented based on Alevin^2^. In the following clustering stage, our framework performs some preprocessing steps (see **Methods**) and uses Louvain algorithm^8^ to cluster the cells based on the initial gene expression profile. Next, in the subpopulation-aware quantification stage, our framework extracts the characteristic of each subpopulation by calculating cluster assigning scores and building the cluster signatures. The gene abundance would also be quantified based on the subpopulation information in this stage. Note that the clustering and the subpopulation-aware quantification stages will be repeated until the clustering results were highly similar between two continuous iterations (Adjust rand index > 0.95).

**Figure 1.**
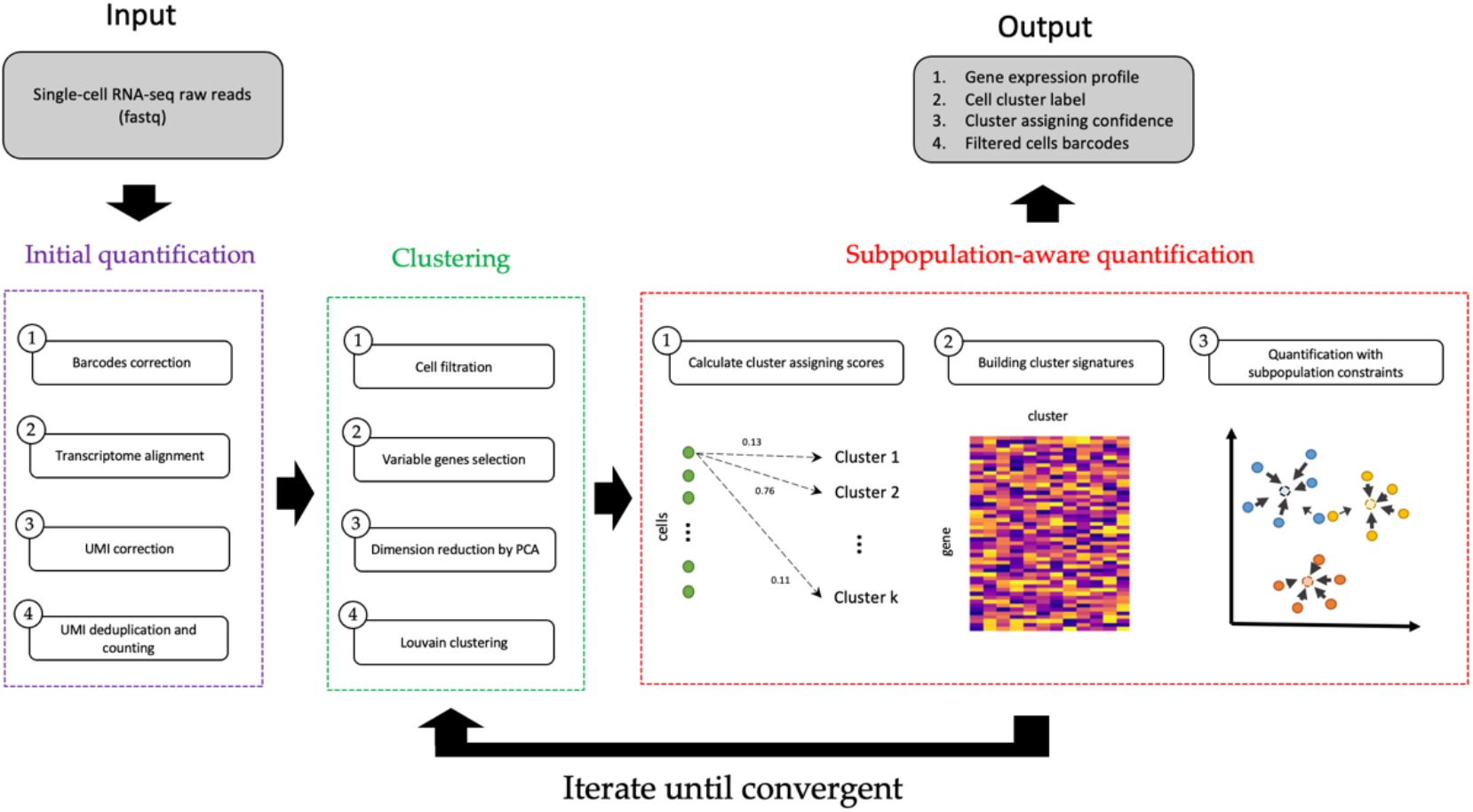
Overview of Quasic. Three stages are contained in the whole pipeline, including (1) Initial quantification, (2) Clustering, and (3) Subpopulation-aware quantification. the clustering and the subpopulation-aware quantification stages will be repeated until the adjusted rand index between two continuous clustering results > 0.95.

### Strengthening of the characteristics and subpopulation structure

In this section, we used the real human PBMCs (peripheral blood mononuclear cells) (see **Methods**) to verify the ability of cell type feature amplification for Quasic. The PBMC dataset contains seven cell types^6^, which were CD4+ T cells, dendritic cells, CD8+ T cells, NK cells, B cells, monocytes, and platelets. In the following paragraph, we would show that the cells in each cell type express significantly different after the subpopulation-aware quantification.

First, we used UMAP^9^ to visualize the association of these PBMC cells. **Fig. 2a-c** showed that compared to other pipelines, our tools aggregated the cells in the same cell type closer to the UMAP space, implying that the features captured by the dimensional reduction methods are more similar for the cells in the same cell type. Second, we also calculated the Pearson correlations between cells and their corresponding cell type’s center using the PBMC markers (see **Fig. 2d**). The results showed that the cell would express the crucial genes more similar to what it was supposed to be due to the properties of cell type. Moreover, we used the package called SAVER^10^, a well-known imputation tool for scRNA-seq published in Nature Methods, to impute the gene expression profiles generated by other quantification pipelines. We compared Quasic to the expression level after the imputation. The results were shown in **Fig. 2e**, showing that despite every quantification tool would express similar to the corresponding cell type after the imputation of SAVER, none of them could amplify the subpopulation structure among the biological data as well as Quasic.

**Figure 2.**
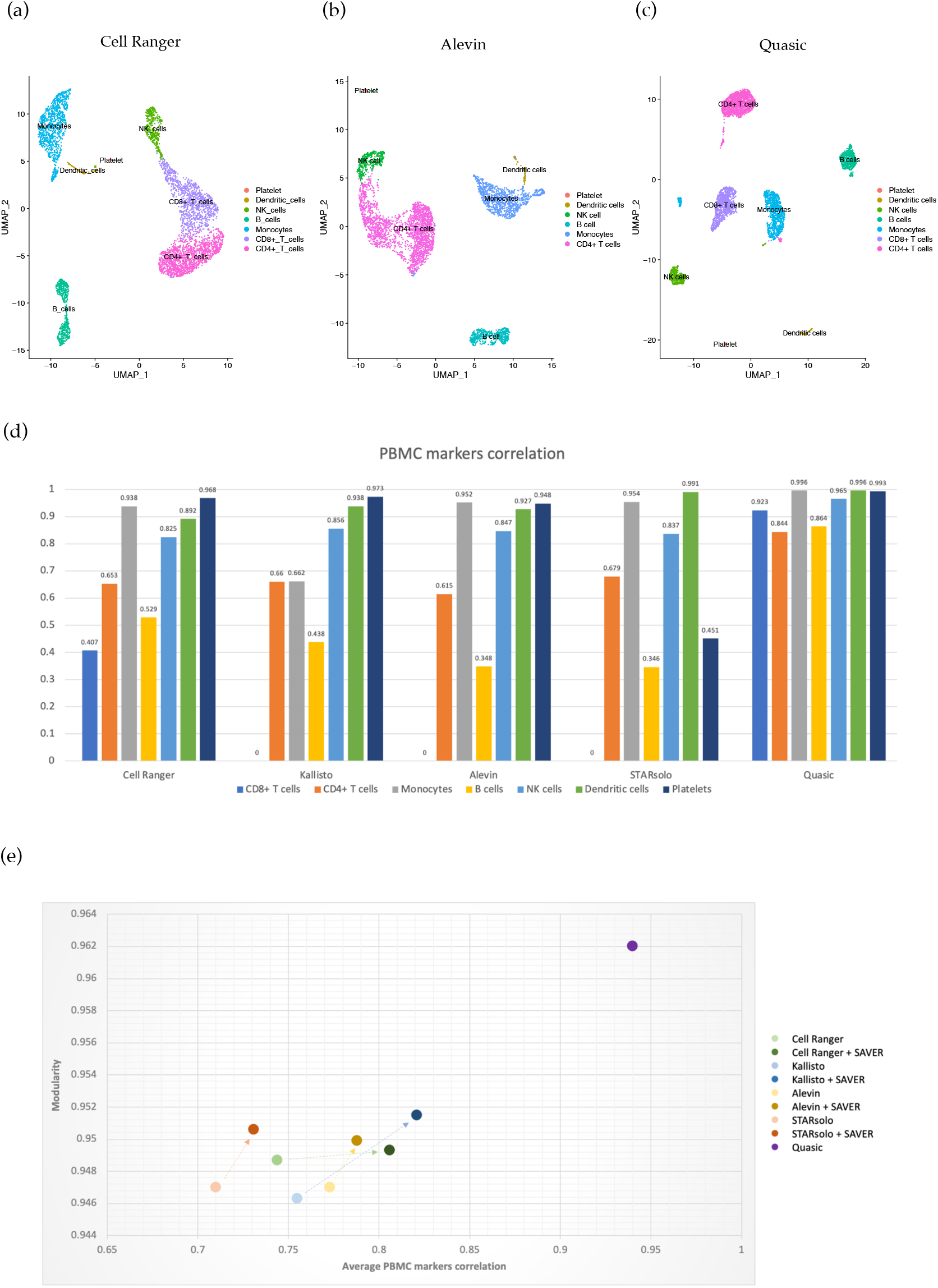
Ability of cell type feature amplification of Quasic. (a)-(c) UMAP visualization of different quantification tools: (a) Cell Ranger, (b) Alevin, (c) Quasic. (d) Pearson correlation between gene expression of cells with corresponding cell type signatures. For each cell type, the mean expression of cells was first calculated, and the Pearson correlations between every cell in this cell type and the corresponding mean expression were further averaged and recorded in the figure. (e) Cell type similarity of different quantification tools before/after the imputation. The modularity used for Louvain optimization was recorded in the figure, and so did the average Pearson correlation of PBMC markers. The results of the same quantification method were colored in similar colors, and the arrow represents the change after the imputation was performed.

Finally, to further examine the effectiveness of downstream analysis for Quasic, we performed the GSEA^11^ (Gene Set Enrichment Analysis), a well-used computational approach that evaluate whether a given gene set shows statistically significant between two or more samples with different biological conditions. We compared the examination of GO (gene ontology) terms on the biological process between different gene expression profiles that were generated from different quantification tools and evaluated the quantities of these GO terms in different q-value thresholds (see **Fig. 3a**). The results showed that in some the cell types (B cells, CD4+ T cells, NK cells), gene expression quantified by Cell Ranger found the minor numbers of GO terms, while Quasic and Cell Ranger used with SAVER were able to help GSEA to find more GO terms within the samples. However, in some cell types, such as Dendritic cells, CD8+ T cells, and Monocytes, GSEA found extremely high amounts of GO terms if SAVER was used for imputation. To check the biological rationality for these GO terms, we used NaviGO^12^ to visualize the relationship between these finding GO terms. **Fig. 3b** showed that for B cells, all three methods found similar groups of GO terms, and most of the GO terms were highly correlated with B cells (phagocytosis recognition^13^, B cells, activation, membrane invagination^14^). In contrast, **Fig. S2** showed that for dendritic cells, Cell Ranger and Quasic found some GO terms that were highly correlated with Dendritic cells. For example, GO terms correlated with major histocompatibility complex (MHC), which is used for presenting antigen-derived peptides in Dendritic cells, were found frequently for the scenario of Quasic, and the correlation between cell adhesion and ATP synthesis toward dendritic cells has also been studied by Harjunpaa et al.^15^ and Wculek et al.^16^. However, the GO terms identified in the condition of Cell Ranger with SAVER were internally irrelevant and poorly correlated with Dendritic cells.

**Figure 3.**
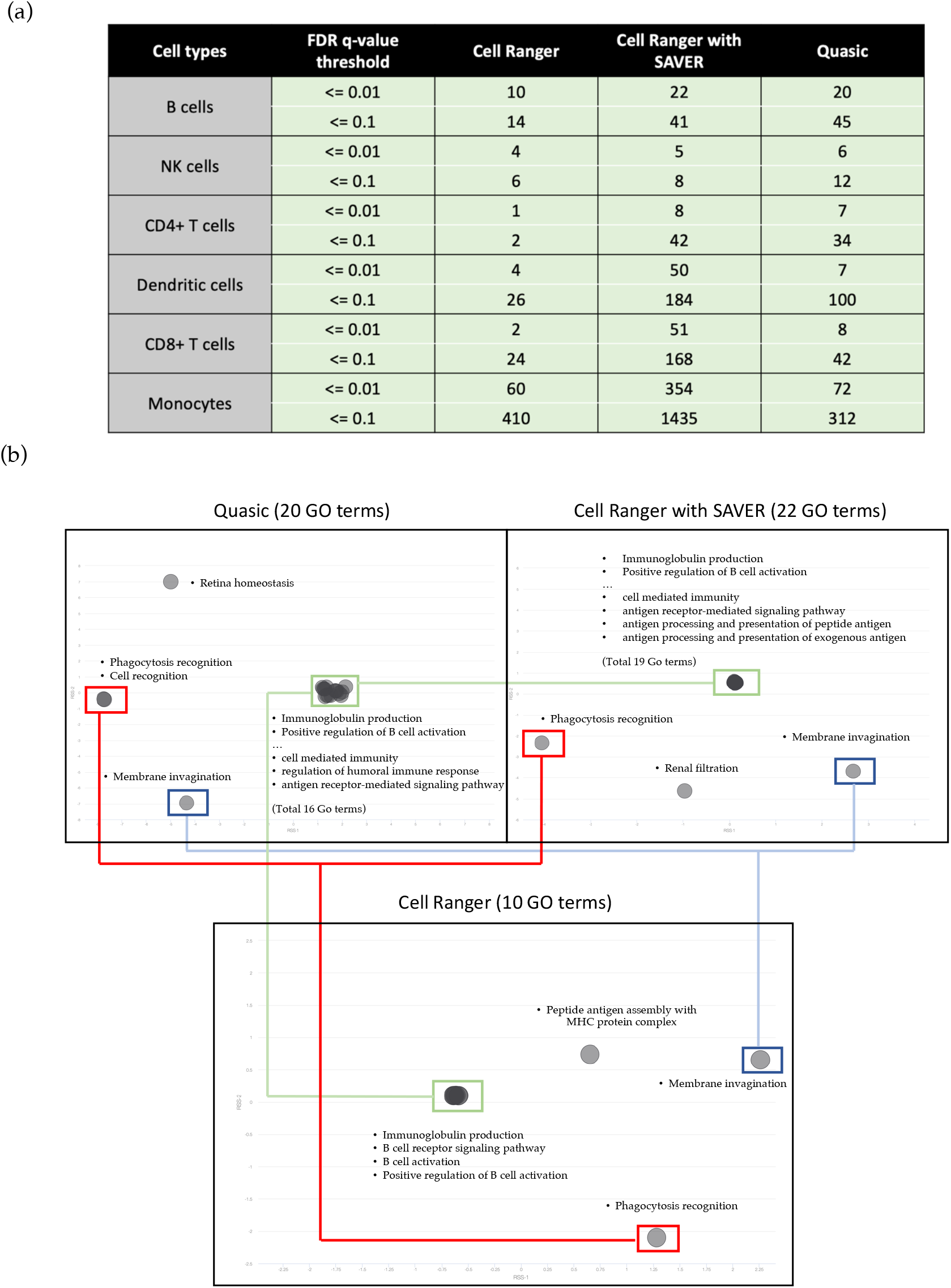
Gene Set Enrichment Analysis results for PBMC datasets. (a) Numbers of GO terms found by multiple quantification conditions in different q-value thresholds. (b) GO terms visualization by NaviGO for B cells. NaviGO would calculate the Resnik Semantic Similarity (RSS) for each GO term pair and use Multidimensional Scaling (MDS) to perform dimension reduction. For each subgraph, the x-axis and y-axis represent the first and second coordinates after MDS. Each black node represents a GO term, and the GO terms with high correlation (which implies their common ancestor were near to themselves) will be close to the figure.

In conclusion, although the imputation tool helps identify more GO terms, it also misrecognizes some GO terms irrelevant to the cell since it amplifies the gene expression level imprecisely. On the other hand, Quasic not only can help the researchers to identify more GO terms than other quantification tools but also avoids the misrecognition that is severely harmful to the researchers to examine the biological condition of their samples.

### Improvements in doublets detection

Droplet-based scRNA-seq platforms often suffer from the “Doublets” issue, which represents two or more cells being considered as a single cell because some GEMs (Gel Beads in Emulsion) mistakenly contain multiple cells during the sequencing library preparation. To avoid affecting the downstream analysis by these doublets, researchers are recommended to identify and filter them before analyzing. In recent years, several tools have been developed to perform doublet detection, such as Scrublet^17^, Scds^18^, and DoubletFinder^19^. In this section, we used the simulation doublets datasets to examine whether Quasic could amplify doublets’ characteristics correctly and benefit doublet detection.

First, we used the single-cell RNA-seq data, which had performed FACS (fluorescence-activated cell sorting) and was purified in cell type to simulate the doublets (see **Methods**). The quantification by Alevin and Quasic were then performed on the simulation data respectively and UMAP was used for visualization. The results in **Fig. 4a-b** clearly showed that Quasic could separate the doublet and the purified cell type cluster. Moreover, since the separation seemed too uncomplicated for the 20% doublets simulation, we added a proportion of ambient RNA to the doublet simulation data. The experiment results for the doublet simulation data with ambient RNA were illustrated in **Fig. 4c-d**, showing that Quasic could still separate the doublet cluster and the purified cell type cluster in the confusing condition. Finally, we used DoubletFinder^19^, a common doublet detection tool for droplet-based scRNA-seq, to examine whether Quasic was a benefit for doublet identification. The experiments were performed on simulation data with different mixing proportions of doublet and ambient RNA (see **Fig. 4e**). The results showed that Quasic is favorable for DoubletFinder recognizing the doublets more accurately than Alevin at all kinds of simulation data, proving that Quasic was able to correctly amplify characteristics of doublets, and was helpful for the doublet detection tool to identify doublets precisely.

**Figure 4.**
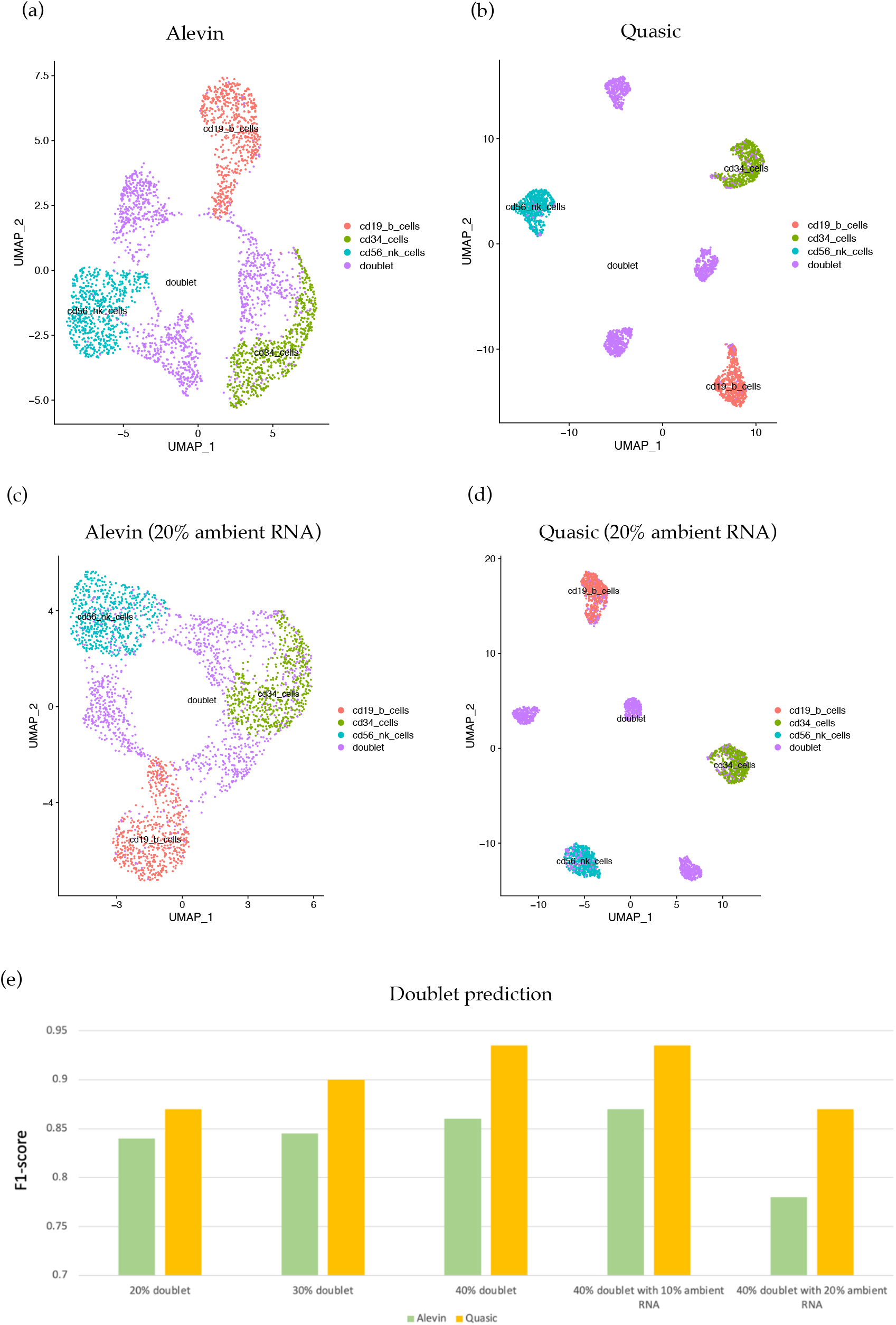
Improvements of doublets detection. (a) – (d) UMAP visualization of different quantification tools for doublet simulation data. (e) F1-score of doublet prediction in different simulation dataset.

### The practicality of identifying rare subpopulation cells in cancer data

In this section, we used a breast cancer scRNA-seq dataset (MCF7, see **Methods**), which contained a rare proportion of cells called pre-adapted cells (PA cells), to prove that Quasic was capable of identifying those rare cells and may be helpful in a practical situation. PA cells are the cells identified in the MCF-7 cell line by Hong et al.^20^, in which the authors examined the transcriptional variability of plastic cells and defined a rare cell subpopulation that displayed increased survival under acute-ET (endocrine therapy) compared with other cells.

To examine whether Quasic was more beneficial in amplifying the features of PA cells, we quantified the scRNA-seq data applied in the study (GSE122743) by Cell Ranger and Quasic, respectively. The UMAP visualization of the quantification results were shown in **Fig. 5a**. It is obvious that while the 81 PA cells, which were identified by Hong et al., were scattered in the UMAP space under the quantification of Cell Ranger, Quasic was able to gather the PA cells into the two small subpopulations. Moreover, considering the biological characteristics of these two subpopulations, it showed that the DEGs (differentially expressed genes) of them are highly overlap with the DEGs found in the original study of PA cells (see **Fig. 5b**). Also, the statistically significant Hallmark gene sets identified by GSEA for Quasic are highly coincide with the gene sets that the original paper had found (**Fig. 5c**).

**Figure 5.**
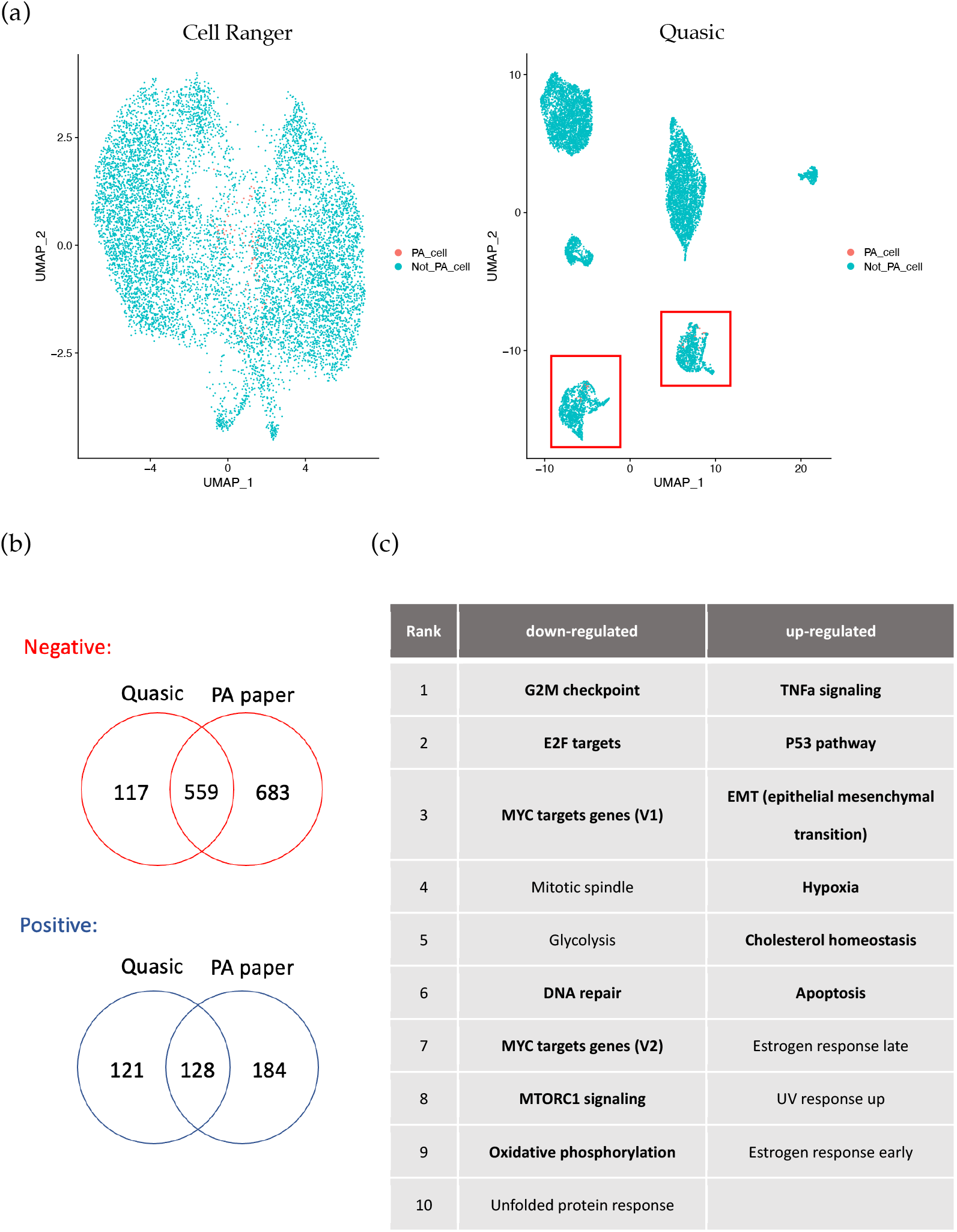
Analysis of MCF-7 dataset and the biological features comparison between two specific subpopulations and PA cells. (a) UMAP visualization of MCF-7 dataset in different quantification tools. The PA cells identified by the previous study were colored in orange, and the two PA aggregated clusters were marked by red frames in the right subgraph. (b) The overlap of positive and negative DEGs between two specific subpopulations an PA cells. (c) The up-regulated genes and down-regulated hallmark gene sets identified by our pipeline quantification, where the letters with boldface represent the hallmark gene sets were also recognized by the original study.

In conclusion, since the biological characteristics between the two subpopulations and the origin identified 81 PA cells were very similar, we considered that these two subpopulations might also contain a high proportion of PA cells, and the potential PA’s features of these two subpopulations could only be recognized under the proposed subpopulation-aware quantification.

## Discussion

Although the above experiments have shown the capability of amplifying the correct gene features and the comparative advantages over the imputation tools for Quasic, the robustness of the proposed subpopulation-aware quantification in some scRNA-seq data with peculiar features still needs to be testified. For example, as mentioned before, the scRNA-seq data usually suffers from dropout events which represent the multiple missing values in the gene expression profiles. According to some previous studies^21^, typical scRNA-seq data can contain up to 90% zero values in the expression matrix. Theoretically, Quasic could recover the expression correctly for these particular circumstances as long as the specific gene features of the cell subpopulation do not lose due to the low sequencing depth, but some related experiments need to be done and testified in the future.

Another notable issue for our proposed framework is the constraint intensity for the subpopulation-aware quantification. Although the current subpopulation constraints have considered the confidence levels of the cluster results for the cells, since cells would be gradually similar to the single cluster signature that had been clustered in, the global intensity of the subpopulation constraint should be set purposely by the users. For example, since the cells in the previous breast cancer experiments (see **Results**) were all from the same cell-line (MCF-7) and the heterogeneities were implicit, only if the global intensity was set at a high value, the aggregation of PA cells could be seen in the UMAP space (see **Fig. S3**). Therefore, the user should decide the global intensity of the subpopulation constraint more purposely since the different constraints would lead to different visible subpopulation structures for the sampled cells and impact the downstream analysis.

Finally, since other scRNA-seq quantification tools have implemented multiple preprocessing strategies before quantification (e.g., different read filtration or UMI deduplication approaches), Quasic should include these preprocessing steps in the future. Moreover, various methods have been developed for clustering scRNA-seq data, and each clustering algorithm has its own advantages in accuracy depending on the attributes of scRNA-seq data. Since the clustering result plays an important role in the correctness of our framework, the effectiveness of different clustering algorithms in alliance with the subpopulation-aware quantification should be further exanimated, and our pipeline should be implemented to allow users to choose their own desired clustering strategies.

## Conclusions

In this study, we proposed a subpopulation-aware quantification pipeline for scRNA-seq data, which aim to strengthen the biological features of subpopulations and favor the downstream analysis. We demonstrated that Quasic can amplify the PBMC markers for each cell type correctly, which not only facilitates cell type identification, but also make the biological pathways found by GSEA more precise and accurate. Furthermore, we used the doublet simulation data to prove that the doublets’ attributes would be recognized clearly through our pipeline, and the features of doublets could also be captured easily by doublet detection tools. In the end, by using Quasic in alliance with differentially expressed gene analysis, we identified more potentially therapeutic resistance cells in the breast cancer data whose characteristics are consistent with previous studies, proving that our pipeline helps discover the rare cell subpopulation in scRNA-seq data.

In conclusion, Quasic can let the gene expression profile of each cell be more consistent with the corresponding subpopulation, making the biological features unique to the subpopulation more apparent and convenient for analysis. Using the proposed quantification pipeline, researchers can effectively extract the desired cell subpopulation information from their sampled cells, enable them to perform cell subpopulation-related studies more accurately.

## Methods

### Materials

#### Real Human 5k Peripheral blood mononuclear cells (PBMCs)

The human PBMC raw reads were downloaded from the <u>10x Genomics website</u>. The data contains about 5025 cells, and 7 different cell types (CD4+ T cells, dendritic cells, CD8+ T cells, NK cells, B cells, monocytes, and platelets) were recognized by Brüning et al.^6^. In our experiments, the raw reads of this PBMC dataset were used as input for different quantification pipelines, and the clustering was further performed by Seurat^22^.

#### Doublet simulation data

We used the PBMC dataset, classified by FACS (fluorescence activated cell sorting), to simulate the heterotypic doublets. The purified cell type of PBMC reads were sequenced from Zheng et al.^1^ and were downloaded from https://www.10xgenomics.com/resources/datasets/. Since some of the cell type datasets were not really purified and may contain other cells with different cell types (which had also been shown in the supplementary of the paper^1^), we only selected CD19+ B cells, CD34 cells and, CD56 NK cells for simulation. Moreover, we filtered the cells which did not express the corresponding markers in the datasets to ensure the correctness of the cell type label.

The doublets were then simulated by mixing reads with two cells in different cell types at a certain proportion (mixing rates were chosen randomly and uniformly between 10% - 90%). Also, 10% / 20% ambient RNA reads, selected from the pool generated by randomly picking 300 cells for each cell type, were added to all cells (including the singlets and doublets) in the simulation dataset.

#### MCF7 with a rare cell subpopulation (pre-adapted cells)

The MCF7 single-cell RNA-seq data was downloaded from Gene Expression Omnibus (GEO) website (https://www.ncbi.nlm.nih.gov/geo/) with the accession number GSM3484478. The PA cells (pre-adapted cells) were identified by Hong et al.^20^, while the barcodes and the differentially expressed genes of PA cells were also obtained from the same paper.

### Subpopulation-aware quantification

As mentioned previously, despite of the correlation between cells and the relevance among or within the cell subpopulation being crucial for downstream analysis, such information is entirely unconsidered in current quantification methods. To model the subpopulation structure within the data, we consider the likelihood model to incorporate a prior that describe the gene expression pattern of each cell type as follows:

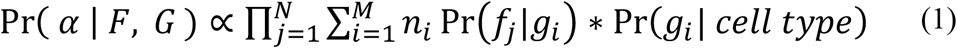

where

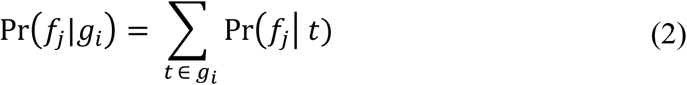

and

The above equations were modified from Salmon^23^ by adding the cell type

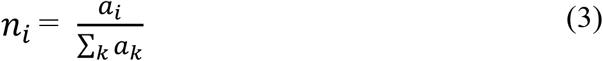

specific term for each cell. For a given fragment set F (including total N fragments obtained after UMI deduplication) and gene set G (total M genes), we are eager to evaluate the gene abundance *α* = {*α*_1,_ *α*_2_, …, *α*_*M*_} (where *α*_*k*_ represents the gene abundance for gene *k*). and the possibility for “fragment *j* is generated by gene *i*” is symbolled as Pr(*f*_*j*_|*g*_*i*_), which is the summation of fragment aligned probabilities of every transcript belonging to gene *i* (**Equation 2**). Moreover, the optimization equation should consider the prior, of which effect is similar to a regularization term to bound the estimation of *g*_*i*_ by the cell type specific signature. Although the prior is unknown, given that cells of the same cell type express similarly, we can still estimate the prior according to the average gene expression among cells that share similar expression profiles. Furthermore, to reduce model complexity, we decided to add the subpopulation-constraint learnt from an additional coarse-grain quantification and clustering step to the original M-step update equation (which had been derived in Salmon^23^) that should achieve the similar bounding effect of the prior for each cell type, see **Equation 4** and **Table 1**.

**Table 1.** Parameters explanation of Equation 4.

| Symbol | Description | Symbol | Description |
| --- | --- | --- | --- |
| $u$ | Iteration index | $C_n$ | Equivalence classes for cell $n$ |
| $i$ | Gene index | $d^j$ | Read counts in equivalence class $j$ |
| $j$ | Equivalence class index | $w_i^j$ | Weight of gene $i$ for equivalence class $j$ |
| $k$ | Cluster of the cell | $\mu_{i,k}$ | Cluster signature of gene $i$ for cluster $k$ |
| $n$ | Cell index | $\lambda_n$ | Intensity of the subpopulation constraint of cell $n$ |
| $\alpha_{i,n}^u$ | Abundance of gene $i$ in cell $n$ for the $u$'th iteration | | |

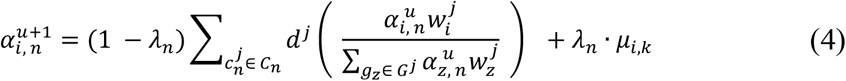

To be more specific, the whole update equation for subpopulation-aware quantification was designed by weighing the gene expression levels calculated by original quantification and the corresponding cluster signature for the cell (*μ*_*i,k*_). Moreover, the weighing was determined by the cell-specific weighting term *λ*_*n*_, which represents the intensity of the subpopulation constraint of cell *n*. In brief, the term *λ*_*n*_ would be higher if the cluster of cell *n* was labeled with strong confidence. Both *μ*_*i,k*_ and *λ*_*n*_ would be introduced in more detail later.

### Cluster assigning score calculation

In the previous section, we mentioned that the parameter *λ*_*n*_ is used for regulating the intensity of the subpopulation constraint for the specific cell *n*. For each cell, the intensity of the subpopulation constraint for the quantification is modeled according to the confidence degree of the cell belonging to its labeled cluster, which means *λ*_*n*,_ ∝ *S*_*n*_, where *S*_*n*_ is the confidence level and called the “cluster assigning score” for cell *n*.

In this study, we calculated cluster assigning scores based on the distance between the cell and the cluster centers in PCA space. The coordinates of each cluster center were calculated by the formula:

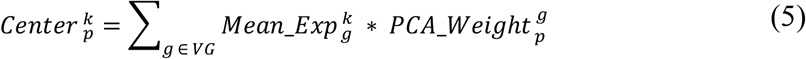

Where *k* and *p* represent the index of cluster and principal component, *g* represents each of the variable genes selected in the previous clustering step. The formula was simply designed as projecting the centers of clusters in variable gene level (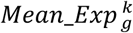, represents the mean expression of variable gene g for cluster k) to the PCA space by multiplying with weights of the specific variable gene for principal components 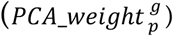. The cluster assigning scores 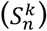 for each cell *n* to each cluster *k* were then calculated:

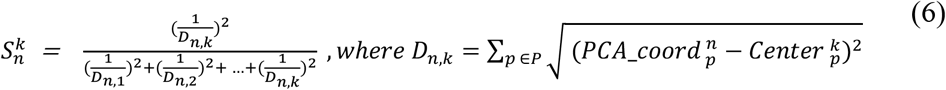

It is worth noticing that the design for the score calculation guarantees the scores of each cluster (*k*) for a cell (*n*) add up to 1 and are inversely proportional to the square distance between the cell and the cluster’s center (*D*_n*k*,_).

Based on the cluster assigning scores, we finally set *λ*_*n*_ = *γ* * *S*_*n*_, where *S*_*n*_ represents the scores for cell n to its assigned cluster labeled according to the former clustering algorithm, and *γ* represents the global intensity of the subpopulation constraint set by users.

### Building of cluster signatures

Generally, the cluster signature for each (*μ*_*i,k*_) was the mean of gene (*i*) expression for the cells in the same cluster (*k*). However, in order to avoid cells that were mis-clustered or low-confidence clustered affecting the correctness of the signatures, we only used the “confidence cells” to calculate the signature. The cell was considered confident if the cluster of maximum clusters assigning scores for this cell was consistent with the cluster that the clustering algorithm labeled, otherwise the cell was considered as an unconfident cell. In conclusion, the cluster signatures are calculated as:

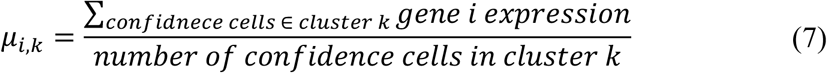

### Low-quality cells and rare gene exclusion

During updating the optimization equation (**Equation 4)**, some cells and genes were excluded from the subpopulation-aware quantification. In terms of cells, cells filtered before the clustering due to their low quality were not included in the subpopulation-aware quantification because they were unlabeled for clustering. Moreover, to avoid amplifying the expression of unimportant genes and make the subpopulation constraint more specific for each cluster, genes which are considered as the “rare genes” (defined as the genes expressed in less than 20% of cells for a cluster) were excluded from the subpopulation-aware quantification.

### Initial quantification

In our framework, we use the methods implemented in Alevin^23^ to perform the preprocessing steps, which includes such as barcodes correction, transcriptome alignment, UMI correction and deduplication. Moreover, we also perform the quantification algorithm of Alevin to estimate the gene expression preliminary before the subpopulation-aware quantification in order to determine the cluster labels for the cells at the first iteration of our framework. All the experiments in the study performed these preprocessing steps and the initial quantification with the default parameters of Alevin.

### Cell Clustering of PBMC, doublet simulation dataset, and MCF7

To calculate the cluster label for each cell (which would be used in subpopulation-aware quantification), the clustering algorithm is included in our framework. However, since scRNA-seq data often contain low-quality cells or numerous univariable genes, multiple preprocessing steps need to be done before clustering. First, cells with detected genes < 20 or mitochondria genes proportion > 10% were excluded for the clustering, and the log normalization is performed and multiplied with a scale factor of 1000. The top 2000 most variable genes are then selected by Seurat^20^ and use for computing the PCs. PCs that accumulated explanation > 90% are further used for clustering and visualization.

In our framework, we use the Louvain algorithm^8^ to perform clustering. Louvain is currently the most popular method for scRNA seq clustering, and is capable of obtaining satisfactory results in most cases with PCA dimension reduction^24^. In our experiments, the parameter in Louvain called “resolution” were set at 0.2 for both PBMC and MCF7 datasets, while was set at 0.1 for doublet simulation dataset.

### Imputation of gene expression profiles by SAVER

SAVER^10^ was downloaded from https://github.com/mohuangx/SAVER. The imputation was executed on the output gene expression profiles of quantification tools directly with default parameters. The gene expression profiles after running SAVER would be further used for PCA dimension reduction and downstream analyses.

### Gene set enrichment analysis

GSEA^11^ software was downloaded from https://www.gsea-msigdb.org/gsea/index.jsp. To analyze of real human PBMC, we first averaged the gene expression level for each cluster and input the target cell type as the phenotype label for GSEA analysis. The permutation type was set to gene set, and the metrics of the ranking gene were set to “log2_Ratio_of_Classes”.

For the MCF-7 dataset, we performed the GSEA pre-ranked analysis. The given PA cells’ barcodes were first labeled as a PA cluster, and the marker genes of this cluster were further obtained by function “Findmarkers” in Seurat^22^ with default parameters. All the upregulated and downregulated genes were used in the GSEA pre-ranked analysis, and their average log fold change calculated during the marker gene finding was used as the ranking score for the corresponding genes.

### GO terms visualization by NaviGO

NaviGO^12^ (https://kiharalab.org/web/navigo/views/goset.php) is a visualization tool used for examining the relationship between GO terms. To perform NaviGO, we first changed the target GO term names, which was the output of GSEA, to corresponding GO term IDs through the Molecular Signatures Database (https://www.gsea-msigdb.org/gsea/msigdb). The “GO Term Similarity and Association Scores” function on the NaviGO website was then performed, and the bubble charts which visualized the relationship of GO terms would be generated.

