## Supplementary for "Reliable and accurate gene expression quantification with subpopulation structure-aware constraints for single-cell RNA sequencing"

### Supplementary figures

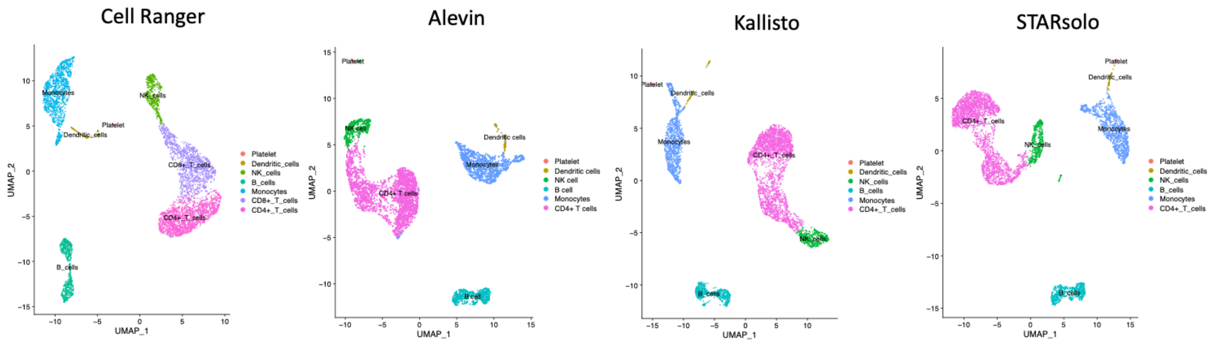


**Figure S1. UMAP visualization of PBMC cells based on different quantification tools.** Cells in different cell types did not separate in the UMAP space due to their ambiguous biological features. The dataset was introduced in **Methods**, and cell types were obtained by calculating the Jaccard similarity scores of each cluster.

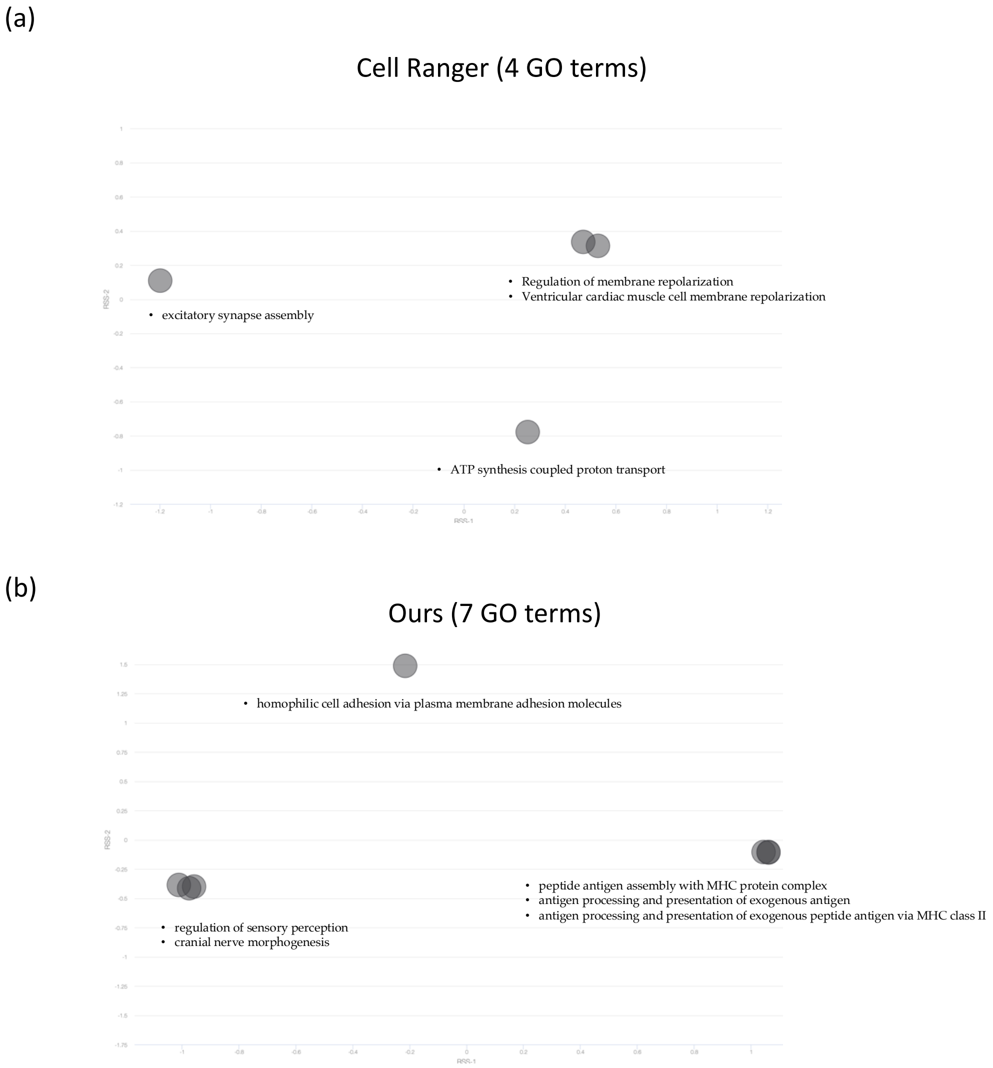

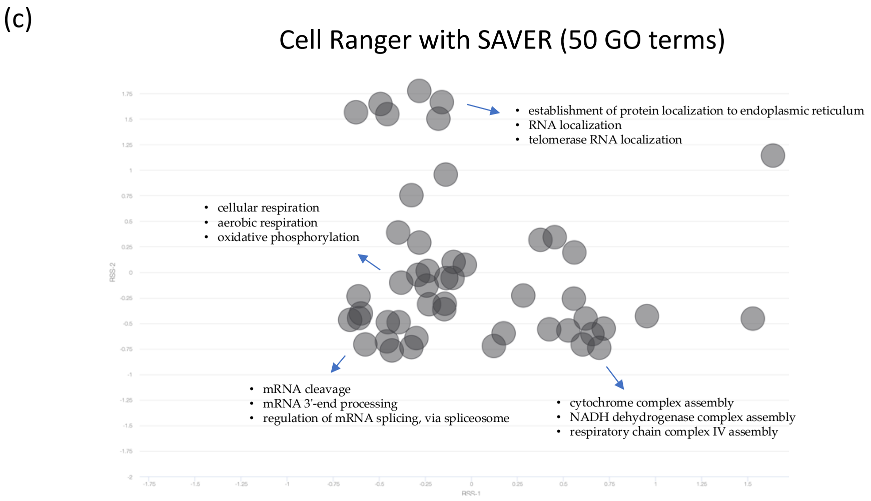


**Figure S2. GO terms visualization by NaviGO for Dendritic cells.** The graph above was generated as same as **Figure 3b**. Each black point represents a GO term, and the GO terms with high correlation (which implies their common ancestor were near to themselves) will be close in the figures.

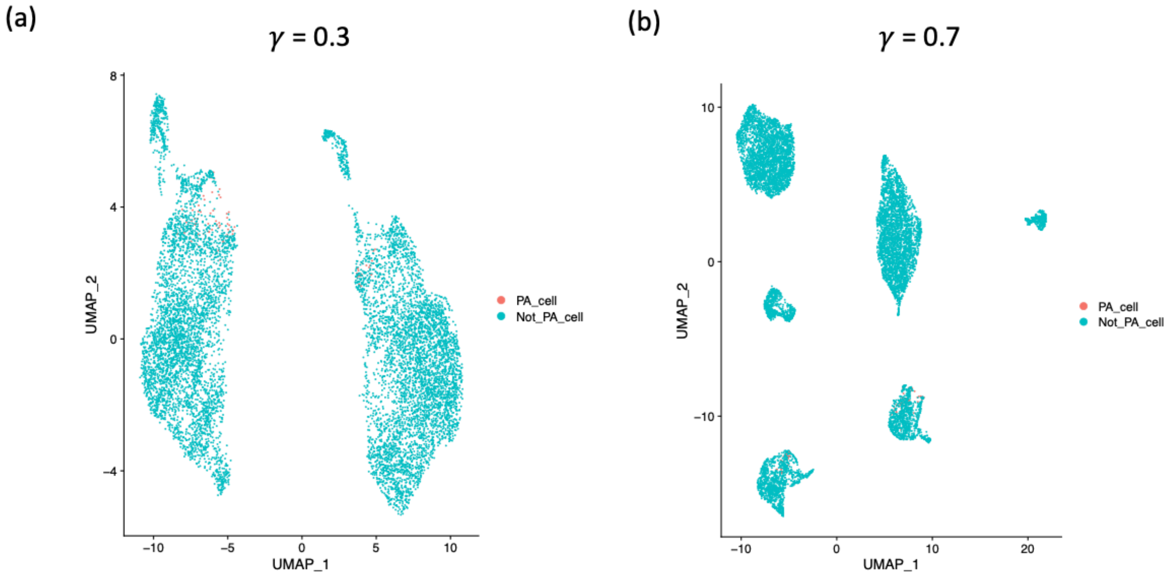


**Figure S3. Quantification results for different intensities of subpopulation-aware constraint in MCF-7.** (a) The global intensity was set at 0.3, and cells were mainly grouped in two clusters based on their different cell cycles. (b) The global intensity was set at 0.7, and PA cells were more gathered in the two small clusters.
